# Methyl substituted *β*–lactam framework based antibiotics and *β*-lactamase inhibitors: Proof of concept by computations

**DOI:** 10.1101/2024.01.14.575563

**Authors:** Vaishali Thakkur, Chandan Kumar Das, Shivani Verma, Suman Saha, Nisanth N. Nair

## Abstract

Increasing the life-time of the acyl-enzyme complex formed between an inhibitor or drug molecule and the *β*-lactamase through chemical modifications of existing drug molecules is an important strategy towards developing inhibitors. In this direction, our group proposed a methyl-substituted *β*-lactam framework for the design of inhibitors for *β*-lactamases (*J. Phys. Chem. B*. **2018**, 122, 4299). This unconventional design was guided by the transition state structure of the deacylation reaction of the acyl-enzyme complex. Here, we present a proof of principle study of this concept through detailed molecular simulations and free energy calculations. In particular, we improve the antimicrobial activity of the first-generation cephalosporin antibiotic, cephalothin, through C6-methylation. The proposed molecule, (6R,7R)-3-(acetyloxymethyl)-6-methyl-8-oxo-7-[(2-thiophen-2-ylacetyl)amino]-5-thia-1-azabicyclo[4.2.0]oct-2-ene-2-carboxylate) slows down the deacylation of the acyl-enzyme complex 10^9^-fold with no apparent effect on its binding to class-C *β*-lactamase and formation of the acyl-enzyme intermediate. The design strategy presented in this work can be further extended to all *β*–lactam antibiotics, like monobactams, carbapenems, cephalosporins, and penicillins.

---

In the early 20^th^ century, the discovery of penicillin^1^ marked an important milestone in the field of medicine, giving rise to the era of antibiotics as a highly effective therapeutic approach. However, the efficacy of antibiotics has been increasingly eroded by the formidable challenge of antibiotic resistance. In their relentless evolutionary quest, bacteria have developed diverse mechanisms to fight antibiotic action.^2,3^ Among these mechanisms, one particularly effective strategy involves the production of *β*–lactamase enzymes, which play a pivotal role in rendering antibiotics ineffective through a complex series of acylation and deacylation reactions, ultimately incapacitating the drug (see Figure 1). Recent years have witnessed a worrisome surge in multi-drug resistant (MDR) bacteria, exhibiting resistance to extended-spectrum *β*–lactam drugs (ESBLs) and even carbapenems, leaving the medical community with alarmingly limited treatment options for these bacterial infections.^4–6^ Consequently, antibiotic resistance has emerged as a global threat, ranking as the leading cause of mortality worldwide for infectious diseases.^7^

**Figure 1:**
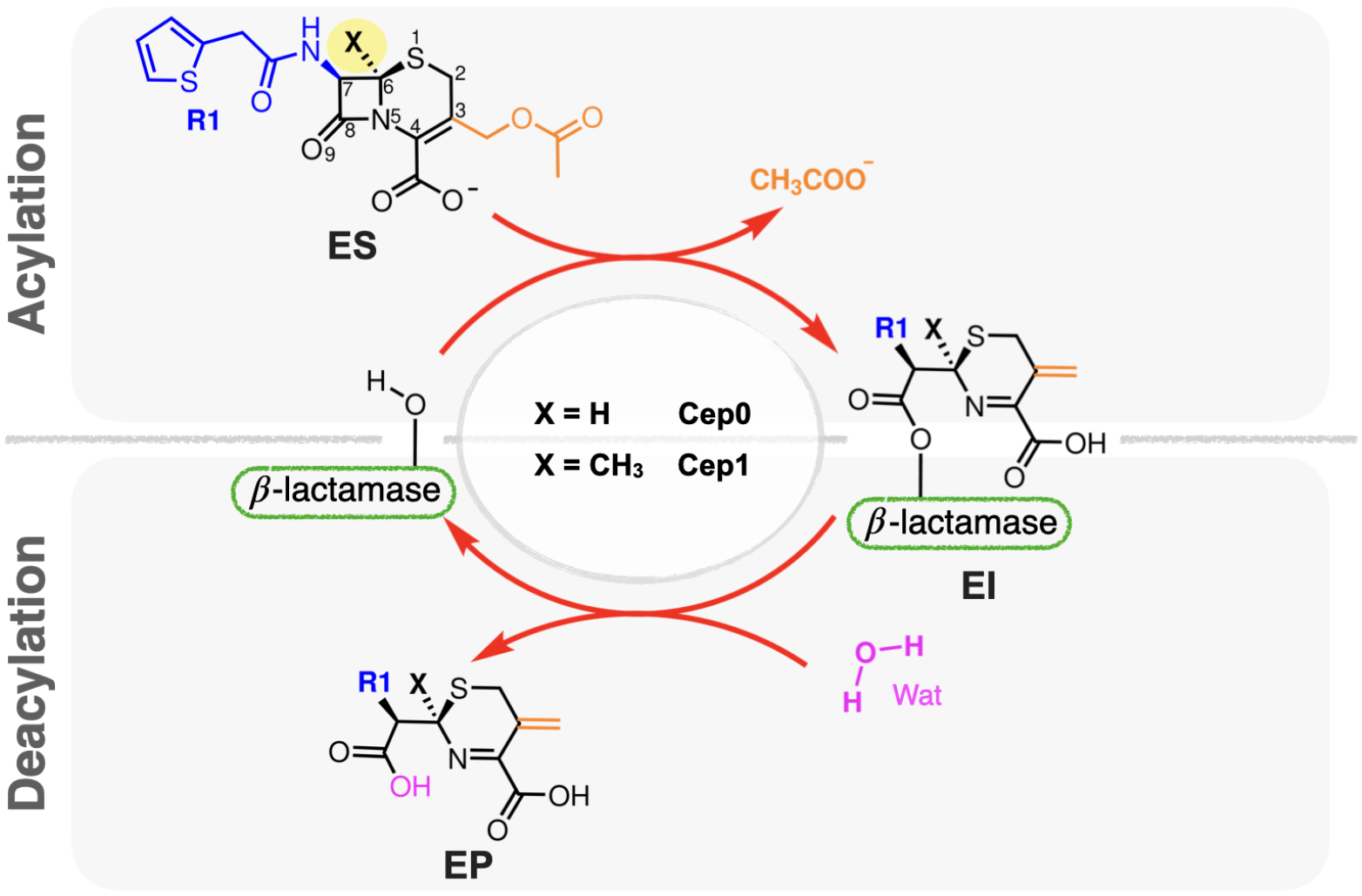
General mechanism of *β*–lactamase catalyzed cephalothin hydrolysis.

In this context, *β*–lactamase inhibitors have emerged as an important resource in the fight against antibiotic resistance. As such, extensive research efforts are currently underway, with a primary focus on unraveling the molecular-level details of antibiotic resistance mechanisms and the development of novel inhibitors.^8–12^ A specific subset of *β*–lactamase enzymes, known as class C *β*–lactamases (CBL) or AmpC *β*–lactamases are specific to gram-negative bacteria^13,14^ and demonstrate resistance against cephalosporins, ceftazidime, and, in certain instances, carbapenems, which are the last resort antibiotics.^15–17^ Notably, monobactams have shown marginal efficacy and form stable covalent complexes, characterized by a slow hydrolytic process.^18^ Our research group has been actively contributing to the understanding of the molecular-level intricacies underlying antibiotic resistance and inhibitor action.^19–26^ Computational investigations have unveiled parallels in the CBL-catalyzed acylation and deacylation processes of cephalothin, a fast hydrolyzing cephalosporin, and aztreonam, a slow hydrolyzing monobactam. During acylation, the role of active site Lys67 as a general base in activating Ser64 for nucleophilic attack has been established, while Tyr150 assumes the role of proton donor.^27^ In the context of deacylation, Tyr150 acts as a general base by facilitating the activation of the water molecule for hydrolysis in both scenarios, with Lys67 contributing as a proton donor to Ser64.^23,24^ Acylation rates for both drugs are similar; however, the hydrolysis of aztreonam is notably slower.^23,24^ Our investigations have suggested a plausible explanation for this discrepancy, linking it to the presence of a methyl group at C4 of aztreonam, which imposes steric hindrance and impedes the incoming hydrolytic water molecule while the transition state of deacylation is formed.^24^ We thus proposed, for the first time, that a methyl substitution at C4 of the *β*–lactam could enhance the antibacterial or inhibitory potential of existing *β*–lactam drugs. Subsequent research has demonstrated the credibility of this logic through the incorporation of C5 substitutions in carbapenems.^28,29^ The resulting drug is reported to show significant activity even against carbapenemases that resist meropenem.^29^ In this work, we further extend our originally proposed idea of a methyl-substituted framework to modify a fast hydrolyzing first-generation cephalosporin, cephalothin.

To this effect, we carried out forcefield-based classical molecular dynamics (MD) simulations to assess the stability of the acyl-enzyme complex with cephalothin (Cep0) and modified cephalothin (Cep1). The free energetics of the hydrolysis reaction were studied using hybrid quantum mechanics/molecular mechanics (QM/MM) MD simulations in conjunction with a highly accurate sampling approach called temperature-accelerated sliced sampling (TASS) technique.^30^ We also compared the free energies of the acylation reaction and drug-binding processes for the two drugs. Additionally, we conducted toxicity and bio-availability analysis of the modified drug to comment on the feasibility of this modification.

The forcefield-based MD simulations were carried out for 1 *μ*s, during which the **EI** covalent complexes, **CBL:Cep0** and **CBL:Cep1**, were found to be structurally stable. The drugs were optimally positioned in the oxyanion hole formed by backbone NH of Ala318 and Ser64. Hydrogen bonding interactions between active site residues, Lys67, Tyr150, Lys312, Asn152, Ala220, Glu272, were conserved in the **CBL:Cep1** complex. Owing to the significance of Lys67-Tyr150 interaction,^22,23,31–33^ we shift our attention to this hydrogen bonding in the conformations sampled during MD simulations. For both complexes, we observed two states, **EI0a**/**EI1a** and **EI0b**/**EI1b**, differing in the Lys67 dihedral conformation. For **CBL:Cep0**, both **EI0a**, and **EI0b** were almost equally populated, while **CBL:Cep1** majorly favoured **EIb**. It can be seen from Figure 2 A and B, that a higher value of Lys67 dihedral is associated with an increase in the distance between Tyr150 and the drug. The presence of methyl group in Cep1 results in a steric clash with Tyr150 at shorter distances, consequently disfavoring **EIa**-like configurations. Additionally, we computed the occupancy of water in the active site of the two complexes and found that **CBL:Cep1** complex has considerably lower water occupancy (Figure 2 C and D). This could be attributed to the C6 methyl, which displaces the active site water. Since the availability of water is necessary for hydrolysis of the **EI** complex, a decrease in water occupancy would lower the probability of hydrolytic attack.

**Figure 2:**
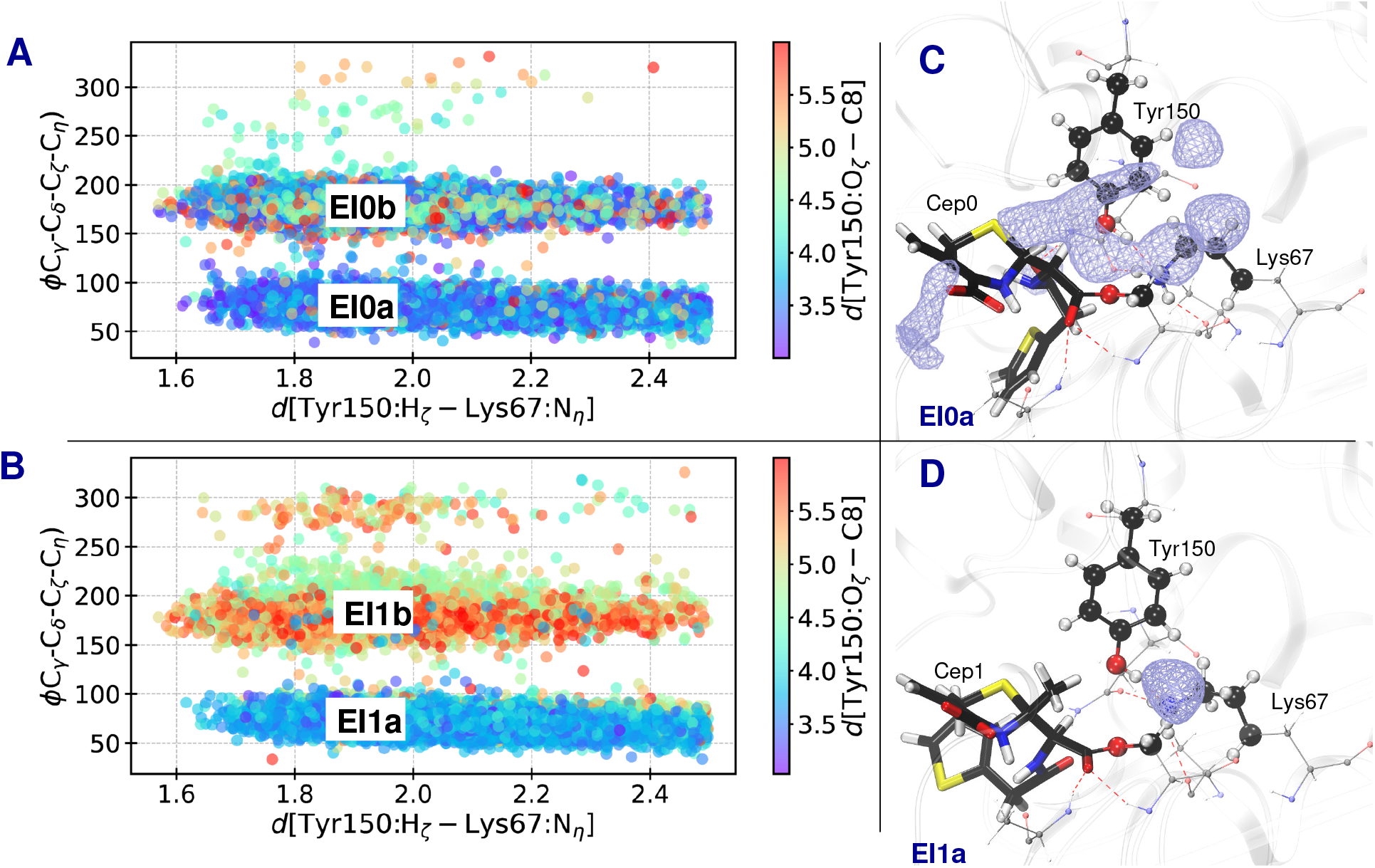
Distance between Tyr150:H_*ζ*_ and Lys67:N_*η*_ plotted against Lys67 dihedral (C_*γ*_– C_*δ*_–C_*ζ*_–N_*η*_) as computed in acyl-enzyme complex of CBL with (A) Cep0 and (B) Cep1. Equilibrated structure of the acyl-enzyme complex **CBL:Cep0** (C) and **CBL:Cep1** (D). The drug is shown in thick stick representation, while the protein residues are shown in ball and stick representation. The pale purple colored mesh shows the region where the average water occupancy is greater than 60%. Atom colors: carbon (black), sulfur (yellow), oxygen (red), hydrogen (white), and nitrogen (blue).

Next, we modeled the hydrolysis of the acyl-enzyme complex. The **CBL:Cep0** hydrolysis is reported to undergo via the formation of a tetrahedral intermediate (**TI)**, which also determines the effective hydrolysis barrier.^23^ We thus studied the mechanism and free energetics of the **EI**→**TI** transformation. In accordance with the previous reports,^23,33,34^ we observed that Lys67:N_*η*_ abstracts a proton from water via Tyr150:O_*ζ*_ thereby initiating the hydrolytic attack on the C8 carbonyl carbon of the drug. Simultaneously, O9 abstracts a proton from N5 to form a tetrahedral intermediate; see Figure 3 A. We computed a barrier of 8 kcal mol^−1^ for **EI0**→**TI0** transformation, indicative of the fast-hydrolysing nature of cephalothin (Figure 3 B, Table 1). On the other hand, the **EI**→**TI1** transformation was found to be significantly slower with a free energy barrier of 20 kcal mol^−1^(Figure 3 C, Table 1). Increase in the hydrolysis barrier of the acyl-enzyme complex from 8 to 20 kcal mol^−1^ translates in to increase in the life-time of the covalent complex by 10^9^ times. We also note that **EI1**→**TI1** transformation observes a late transition state owing to the less stable product **TI1**. This is accompanied by a low barrier for the reverse transformation, **TI1**→**EI1**, suggesting a possibility of regeneration of the acylated intermediate. Analysis of the deacylation transition states, **TSd:Cep0** and **TSd:Cep1**, revealed that the presence of the outward protruding C6-methyl in Cep1 sterically obstructs the incoming hydrolytic water molecule, resulting in a high energy transition state; see Figure 3 C and D.

**Figure 3:**
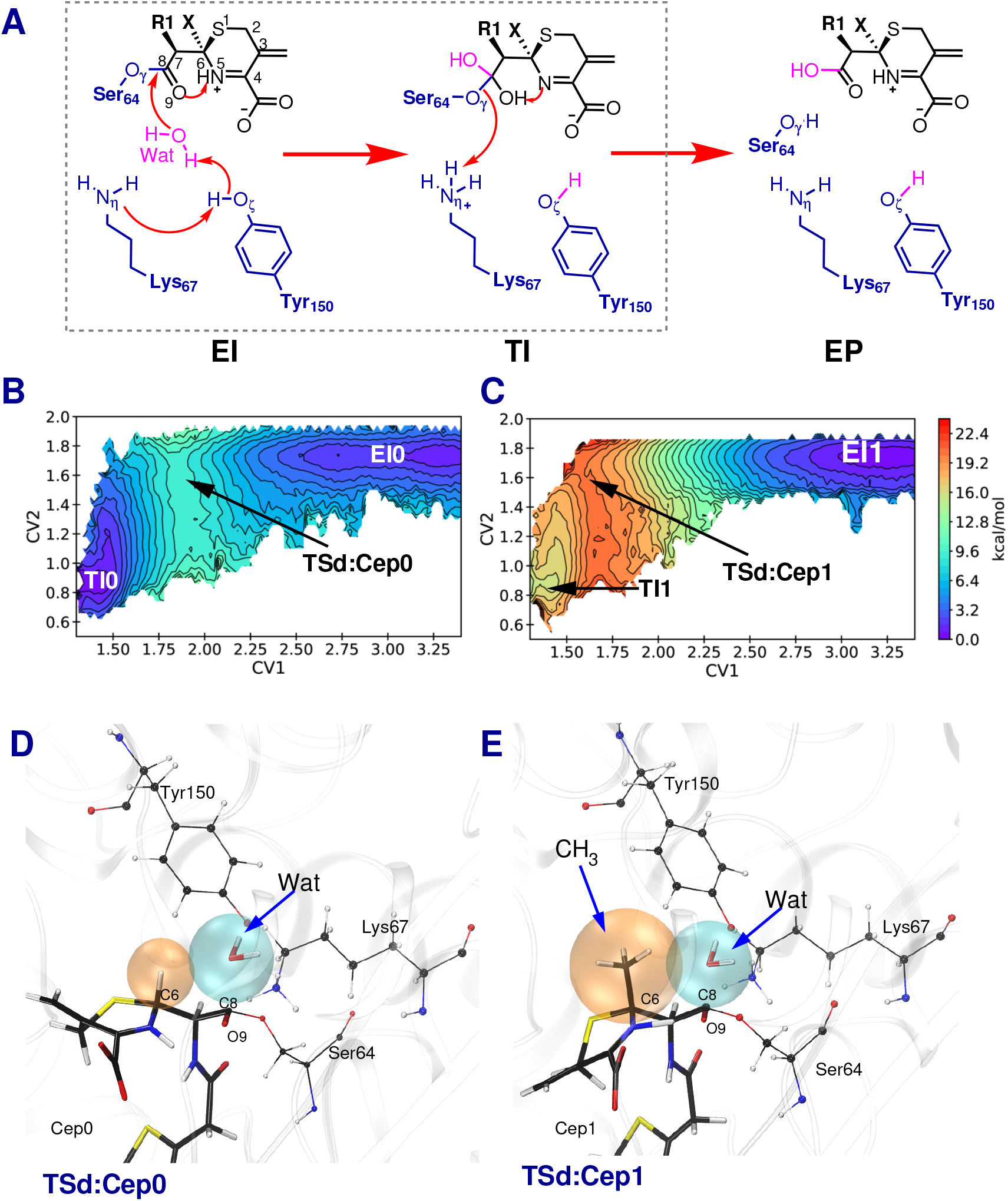
(A) The observed mechanism of deacylation of **CBL:Cep0** (X = H) and **CBL:Cep1** (X = CH_3_). Free energy surfaces for deacylation of **CBL:Cep0** (C) and **CBL:Cep1** (D) are shown as a function of attacking water oxygen (Wat:O) to C8 distance (CV1), and the coordination number of Wat:O with the two H atoms of Wat (CV2). The transition state structures for the deacylation of **CBL:Cep0** (D) and **CBL:Cep1** (E) are also shown. Transparent spheres in (D) and (E) are drawn centered at C6-proton, C6-methyl carbon, and attacking water oxygen with their radii equal to van der Waals radius of methyl (2.0 Å), proton (1.2 Å) and oxygen (1.5 Å), respectively, in order to visually compare the steric clashes.

**Table 1:** Comparison of computed free energy barriers of CBL-catalyzed acylation and deacylation of Cep0 and Cep1.

| Reaction | $\Delta F^\ddagger$ (kcal mol<sup>-1</sup>) | |
| --- | --- | --- |
|  | Cep0 | Cep1 |
| Acylation | 17 | 17 |
| Deacylation | 8 | 20 |

From our simulations, we conclude that the C6-substituted cephalothin influences (a) the occupancy of hydrolytic water in the active site and (b) the organization of active site residues. Both of these factors contribute to the high energy barrier and destabilization of the transition state. However, it is crucial to highlight here that the C6-methyl substitution in Cep1 could slow down the acylation reaction and destabilize the drug-binding interactions as well. In fact, acyl-enzyme complex formation is reportedly the rate-determining step for CBL-catalyzed hydrolysis of cephalothin.^23^ Thus, it is crucial to study the effect of methyl substitution on these processes before we can comment on the inhibitory activity of the modified drug. Consequently, we modeled the formation of acyl-enzyme intermediate (**ES**→**EI**). In line with our previous report,^27^ Lys67 abstracts a proton from Ser64 to initiate the nucleophilic attack of Ser64:O_*γ*_ on C8, leading to the *β*–lactam ring opening. Simultaneously, the conjugation between N5 and the C3–C4 double bond aids in the expulsion of the CH_3_COO^−^ group. This is followed by a proton transfer from Lys67 to the carboxylate group of the drug via Tyr150, see Figure 4 A. The computed acylation-free energy barriers for **CBL:Cep0** and **CBL:Cep1** were found to be the same (∼ 17 kcal mol^−1^); see Figure 4 B and C, and Table 1. This can be explained by the negligible increase in the steric clash in the acylation transition state due to C6-methyl (Figure 4 D and E). Thus, methyl substitution at the C6 position does not affect the nucleophilic attack of Ser64:O_*γ*_ on C8.

**Figure 4:**
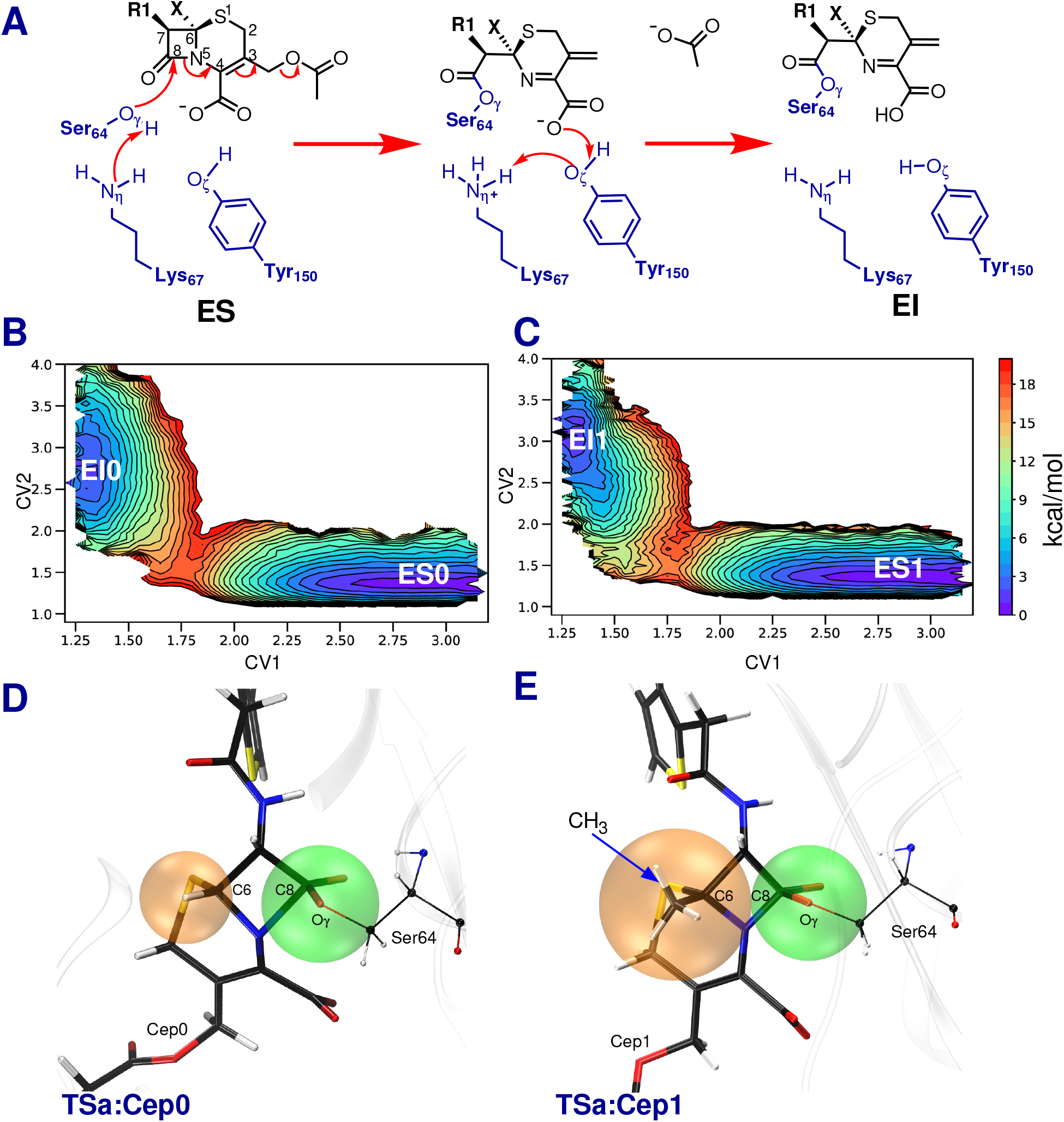
(A) The observed mechanism of acylation is shown for Cep0 (X = H) and Cep1 (X = CH_3_). Free energy surfaces for the acylation reaction (**ES**→**EI**) undergone by Cep0 (B) and Cep1 (C) are given. The transition state structures for these reactions are in D and E, respectively. Transparent spheres in (D) and (E) are drawn centered at C6-proton, C6-methyl carbon, and Ser64:O_*γ*_ with their van der Waals radii as in Figure 3.

Next, we investigate the effect of the modification on the drug-binding affinity in the formation of Michaelis complex. We computed the free energy difference (ΔΔ*F*) of binding between the two drugs to be ∼ 0.1 kcal mol^−1^. Similar binding affinities for the drugs suggest that the introduced modification does not interfere with any protein-drug interactions within the active site. Finally, we assess the toxicity and bio-availability of the potent drug molecule. The predicted median lethal dose (LD50) value for Cep1 is 10, 000 mg kg^−1^ which is comparable to ∼ 10, 000 mg kg^−1^ for Cep0,^35^ indicating that Cep1 has low toxicity.^36,37^ Cep1 was also found inactive for carcinogenicity, immunotoxicity, heptatoxicity, and cytotoxicity. The molecule was also predicted to have acceptable bio-availability with flexibility, lipophilicity, saturation, solubility, and molecular weight all falling within the acceptable range.^38^

Our computations have revealed that the C6-methyl group in Cep1 displaces active site water, thereby slowing down the hydrolysis of the acyl-enzyme complex by nearly nine orders of magnitude. As a result of the increased lifetime of the acyl-enzyme complex, the inhibitory nature of Cep1 turns out to be significantly higher than Cep0. Interestingly, acylation and drug-binding processes are unaffected by this modification. The Cep1 is also not toxic. These findings are especially significant to developing novel inhibitors as most commonly used antibiotics have a *β*–lactam core. Methylation of the *β*–lactam core in the existing antibiotics, as in Figure 5, should be explored further. The synthesis of such molecules needs to be investigated in the future.

**Figure 5:**
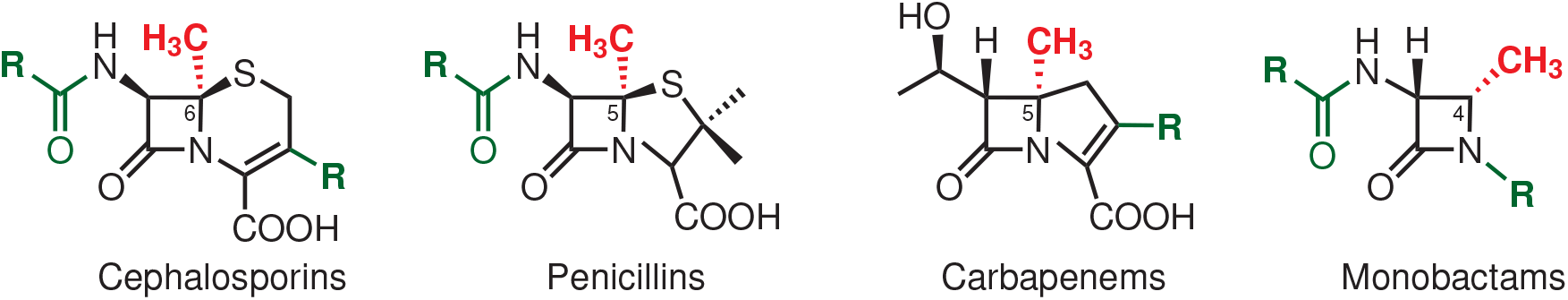
Proposed modifications to the *β*-lactam core of existing *β*-lactams.

## Acknowledgement

Computational resources were provided by the HPC facility and the Param Sanganak (National Supercomputing mission) facility at IIT Kanpur. VT thanks IITK for the PhD fellowship.

